# Rationally designed *mariner* vectors to allow functional genomic analysis of *Actinobacillus pleuropneumoniae* and other bacteria by transposon-directed insertion-site sequencing (TraDIS)

**DOI:** 10.1101/433086

**Authors:** Janine T Bossé, Yanwen Li, Leon G. Leanse, Liqing Zhou, Roy R Chaudhuri, Sarah E Peters, Jinhong Wang, Gareth A. Maglennon, Matthew TG Holden, Duncan J Maskell, Alexander W Tucker, Brendan W Wren, Andrew N Rycroft, Paul R Langford, on behalf of the BRaDP1T consortium

## Abstract

Transposon Directed Insertion Sequencing (TraDIS) is a high-throughput method for mapping insertion sites in large libraries of transposon mutants. The *Himar1* (*mariner*) transposon is ideal for generating near-saturating mutant libraries, especially in AT-rich chromosomes, as the requirement for integration is a TA dinucleotide. In this study, we generated two novel *mariner* vectors, pTsodCPC9 and pTlacPC9 (differing only in the promoter driving expression of the transposase gene), in order to facilitate TraDIS identification of conditionally essential genes in *Actinobacillus pleuropneumoniae* and other bacteria. Using the pTlacPC9 vector, we have generated, for the first time, saturating *mariner* mutant libraries in both *A. pleuropneumoniae* and *Pasteurella multocida* that showed a near random distribution of insertions around the respective chromosomes. A preliminary screen of 5000 mutants each identified 8 and 15 genes, respectively, that are required for growth under anaerobic conditions.

## Importance

Comprehensive identification of conditionally essential genes requires efficient tools for generating high-density transposon libraries that, ideally, can be analysed using next-generation sequencing methods. The *mariner* transposon has been used for mutagenesis of a wide variety of bacteria, however plasmids for delivery of this transposon do not necessarily work well in all bacteria. In particular, there are limited tools for functional genomic analysis of Pasteurellaceae species of major veterinary importance, such as swine and cattle pathogens, *Actinobacillus pleuropneumoniae* and *Pasteurella multocida*, respectively. Here, we have developed plasmids that allow delivery of mariner and the production of genome saturated mutant libraries for both of these pathogens, but which should also be applicable to a wider range of bacteria. High-throughput screening of the generated libraries will identify mutants required for growth under different conditions, including *in vivo*, highlighting key virulence factors and pathways that can be exploited for development of novel therapeutics and vaccines.

## Introduction

*Actinobacillus pleuropneumoniae*, a member of the *Pasteurellaceae*, is the causative agent of porcine pleuropneumonia, a highly contagious, often fatal, respiratory disease that causes considerable economic losses to the swine industry worldwide (1). Certain virulence factors have been shown to have specific roles in the pathogenesis of *A. pleuropneumoniae* infection including RTX toxins, capsule, lipopolysaccharide and various outer membrane proteins (2, 3). In addition, two signature-tagged mutagenesis (STM) studies have identified a large number of genes that contribute to the ability of *A. pleuropneumoniae* to survive and cause disease in pigs, though neither screen was saturating (4, 5).

For more than two decades, Tn*10* has been the transposon of choice for generating libraries of random mutants of *A. pleuropneumoniae* for STM and other studies (4–9). However, Tn*10* has an insertion site preference for GCTNAGC (10), and different insertional hotspots were reported in both *A. pleuropneumoniae* STM studies (4, 5, 11), limiting the usefulness of this transposon for creating a fully saturating library. Clearly, a more random transposon mutagenesis system in *A. pleuropneumonia*e is required in order to allow genome-wide analysis of fitness using high-throughput sequencing methods such as Transposon Directed Insertion-site Sequencing (TraDIS) that not only precisely map, but also quantitatively measure the relative abundance of each transposon insertion in a pool of mutants (12–14).

The *Himar1* (*mariner*) transposon, originally isolated from the horn fly, *Haematobia irritans*, has been shown to insert randomly into the chromosomes of a wide range of bacteria with a dinucleotide target of “TA” (15). In particular, *mariner* has been used for mutagenesis of *Haemophilus influenza* (12, 16), another member of the *Pasteurellaceae*, with a comparatively AT-rich genome, like *A. pleuropneumoniae*.

The aim of this study was to develop a *Himar1 mariner* mutagenesis system for use in *A. pleuropneumoniae*, and other bacteria such as *Pasteurella multocida*, that would be amenable to TraDIS (14).

## Results

### Construction and evaluation of *mariner* vectors

Two novel mobilizable *mariner* delivery vectors, pTsodCPC9 and pTlacPC9 (Figure 1), differing only in the promoter driving expression of the C9 transposase gene, were successfully generated using pGEM-T as the vector backbone. Conjugal transfer into *A. pleuropneumoniae* MIDG2331 from the DAP-dependent MFD*pir*, achieved for both vectors, confirmed that the *oriT* and *traJ* sequences were sufficient to facilitate mobilization. Comparison of results for *A. pleuropneumoniae* indicated that although the *sodC* promoter is constitutively expressed (17), IPTG induction of expression of the C9 transposase from the *lac* promoter resulted in higher frequencies of transposition (10^−6^-10^−8^ compared to 10^−7^-10^−10^). Furthermore, similar frequencies of transposition were found in the genome of *P. multocida* MIDG3277 (10^−6^-10^−8^) using pTlacPC9, whereas no transposants were recovered using pTsodCPC9 in this species. Therefore, the pTlacPC9 vector was used for generation of libraries in both *A. pleuropneumoniae* and *P. multocida*.

**Figure 1.**
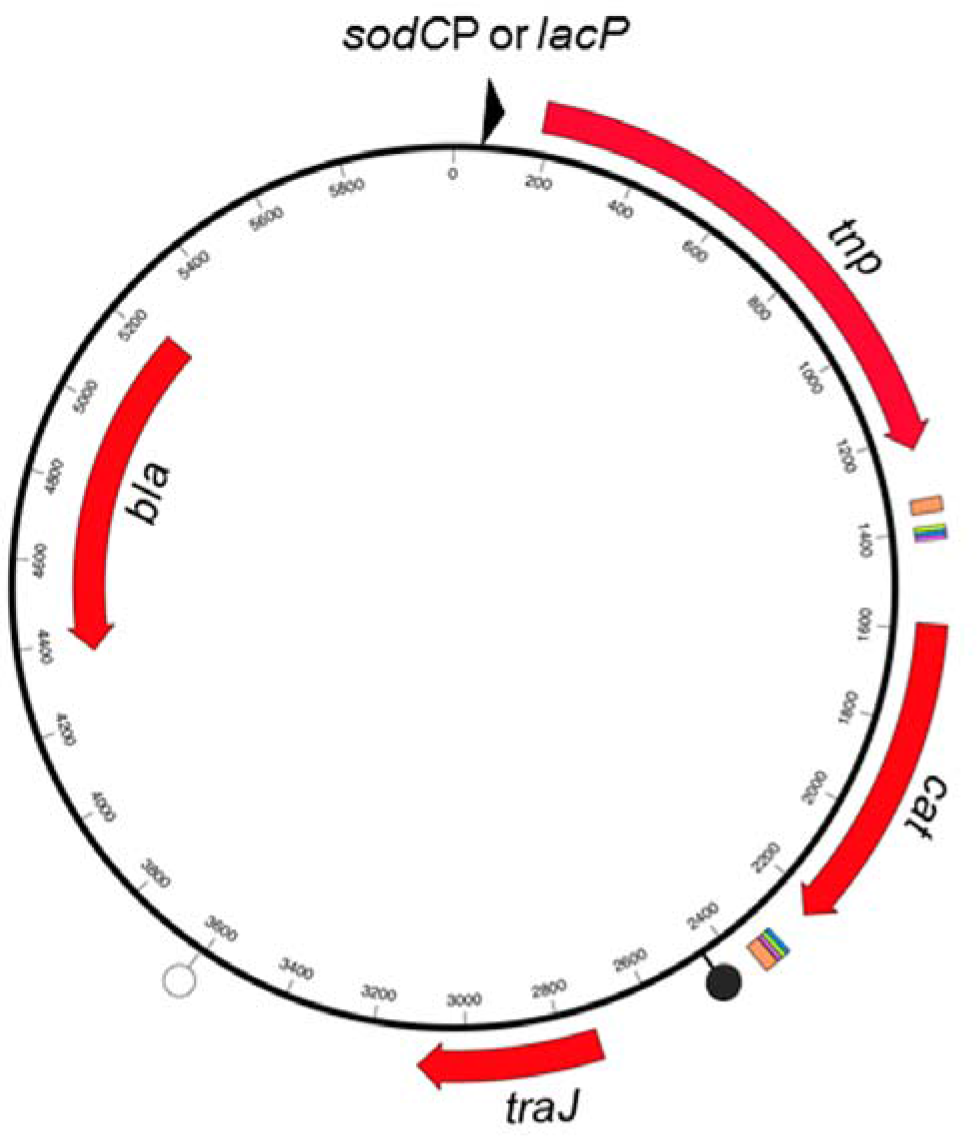
General map of the *mariner* vectors, pTsodCPC9 and pTlacPC9, differing only in the promoter for the *tnp* gene. Genes *tnp* (*mariner* transposase C9 mutant), *cat* (chloramphenicol resistance), *traJ* (plasmid transfer gene), and *bla* (ampicillin resistance) are indicated by the appropriately labelled solid red arrows. The origin of plasmid transfer (*oriT*) is shown as a filled lollipop 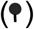, and the origin of plasmid replication (*colE1*) as a hollow lollipop (⫯). Coloured blocks flanking the *cat* gene indicate the locations of the *Himar1* inverted repeats (thick orange), and paired copies of DNA uptake sequences for *Neisseria* spp. (thin green), *H. influenzae* (thin pink) and *A. pleuropneumoniae* (thin blue). Arrowhead upstream of C9 *tnp* gene indicates the promoter sequences for either the *A. pleuropneumoniae sodC* or the *E. coli lac* gene, depending on the vector (pTsodCPC9 and pTlacPC9, respectively).

Colony PCR revealed that initial transconjugants retained extrachromosomal plasmid, as indicated by amplification of both the chloramphenicol (Cm) cassette and the *oriT/traJ* sequence (data not shown). Following subculture on selective agar, only the Cm cassette could be amplified from selected mutants, indicating loss of the plasmid and integration of the transposon into the chromosome. Southern Blot (not shown) and linker-PCR (Figure 2) confirmed single insertions in different locations in 12 randomly selected mutants. Insertions were stable in the absence of selection for 20 generations.

**Figure 2.**
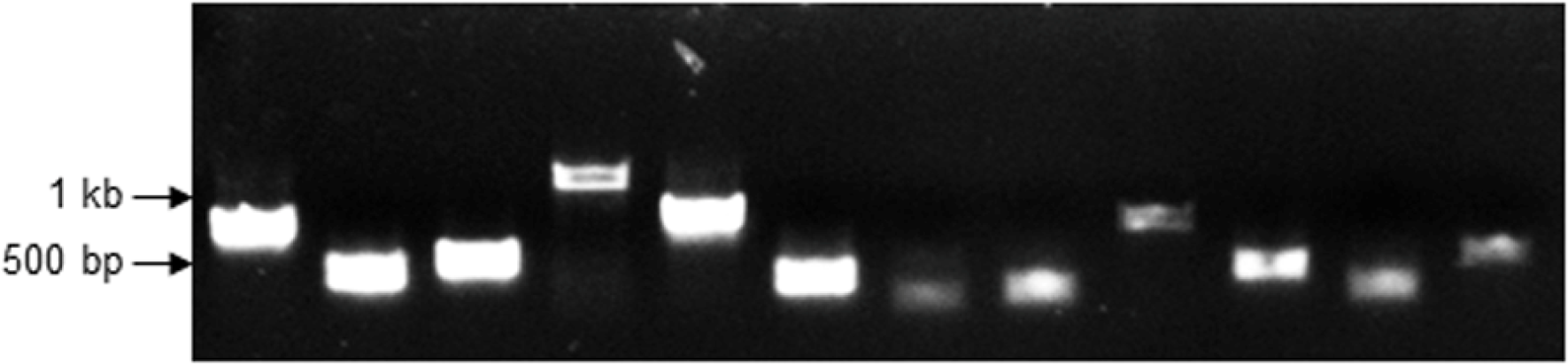
Linker PCR products for 12 randomly selected *A. pleuropneumoniae* mutants. Sequences flanking the *mariner* insertions were amplified from AluI-digested linker-ligated DNA fragments using primers L-PCR-C and IR-Left_out (for amplification of the left flank).

### TraDIS analysis of the *A. pleuropneumoniae* and *P. multocida mariner* libraries

Analysis of linker-PCR products generated from DNA (+/− ISceI digestion) from the pooled *A. pleuropneumoniae mariner* library containing ≥ 43,000 transconjugants showed a dominant plasmid-specific band only in the sample that was not treated with ISceI (Figure 3). Both samples showed a strongly stained smear of DNA ranging from 100 to 500 bp in size, indicating a good distribution of insertions in the library. Subsequent TraDIS analysis of the ISceI-digested *A. pleuropneumoniae* library DNA generated 16,565,883 sequence reads from the 5’ end of the *mariner* transposon. Alignment of the reads against the closed MIDG2331 genome (18) identified 56,664 unique insertion sites in total, with near random distribution around the entire genome (Figure 4A).

**Figure 3.**
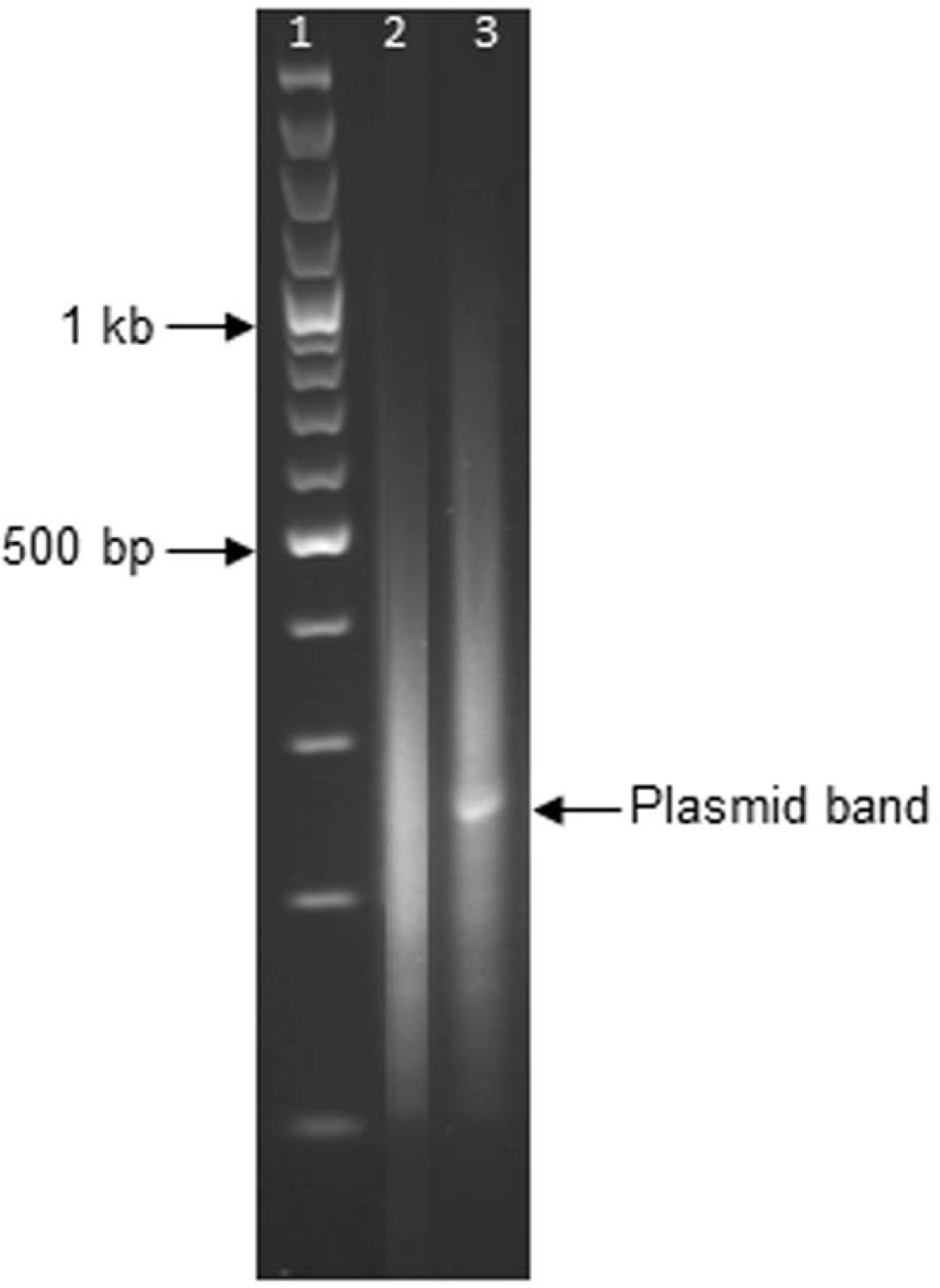
Comparison of linker PCR-products generated from pooled *A. pleuropneumoniae* library genomic DNA +/− ISceI digestion. Lanes: 1) 100 bp ladder; 2) ISceI-treated sample; 3) untreated sample. Amplification products were generated for the left flanking sequences, as in Figure 2. The dominant plasmid band in the untreated sample is indicated.

**Figure 4.**
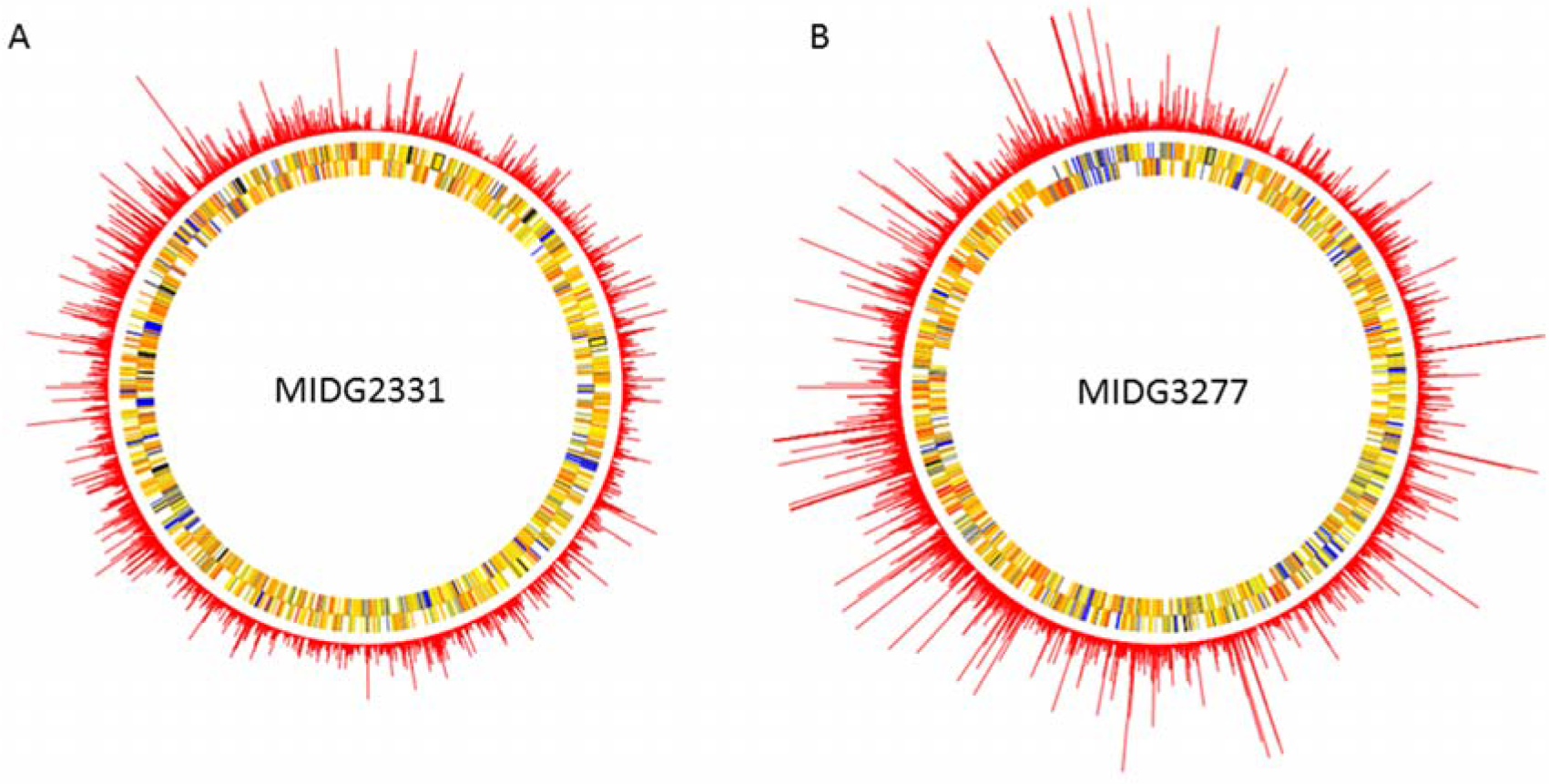
Distribution of *mariner* insertions identified in the *A. pleuropneumoniae* and *P. multocida* genomes. TraDIS reads for the respective pooled libraries were mapped to A) the complete genome of *A. pleuropneumoniae* MIDG2331 (accession number LN908249); and B) the draft genome of *P. multocida* MIDG3277 (accession number ERZ681052). Each spike plotted from the chromosome represents a single insertion site, with the length of each spike proportional to the number of mapped sequence reads from that insertion site. 56,664 unique insertion sites were identified in the *A. pleuropneumoniae* library, and 147,613 in *the P. multocida* library. In the *P. multocida* dataset, there were several insertion sites with large numbers of mapped reads. To enable insertions with fewer reads to be seen clearly, read coverage has been capped to a maximum of 50,000 in B (the true maximum coverage at an insertion site was 136,062 reads).

For analysis of the *P. multocida mariner* library, it was first necessary to generate a draft genome sequence for the isolate used, MIDG3277. Sequencing on an Illumina HiSeq 2000 yielded 2285355 pairs of 100 bp sequence reads, which were assembled into 160 contigs, which were ordered based on a comparison with the genome of *P. multocida* PM70 (GenBank accession AE004439; Figure 5). The assembly had a total size of 2,562,694 bp, with an n50 of 77,141 bp. Annotation of the contigs identified 2490 coding sequences, together with 35 tRNA genes and 4 rRNA genes.

**Figure 5.**
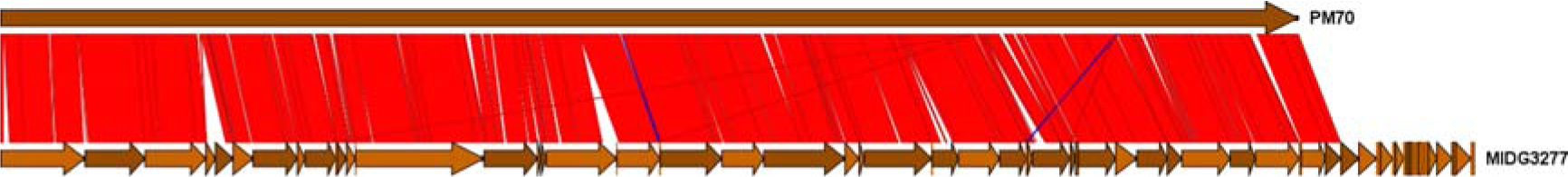
A comparison of the *P. multocida* MIDG3277 draft assembly with the complete genome sequence of *P. multocida* PM70. Arrows indicate the position and orientation of the contigs. Red blocks indicate matches in the same orientation, blue blocks indicate matches in the reverse orientation. The assembled contigs of MIDG3277 were ordered based on this comparison, which was performed using NUCmer 4.0 (43).

TraDIS analysis of the ISceI-digested *P. multocida* library DNA generated 46,809,272 sequence reads, from the 5’ end of the *mariner* transposon. Alignment of the reads with the draft genome sequence of MIDG3277 identified 147,613 unique insertion sites, with distribution around the entire genome (Figure 4B).

### Preliminary screen for anaerobic mutants

Of approximately 5000 mutants each of *A. pleuropneumoniae* and *P. multocida* (from 4 different matings in both cases) screened for the presence of insertions in genes required for anaerobic growth, 19 and 15 mutants, respectively, were identified that failed to grow. Linker-PCR produced amplicons of different sizes that, when sequenced, mapped to unique TA dinucleotides in 8 different genes in the *A. pleuropneumoniae* MIDG2331 genome (Table 2), and 15 different sites (some intergenic) in the *P. multocida* MIDG3277 genome (Table 3).

**Table 1.**
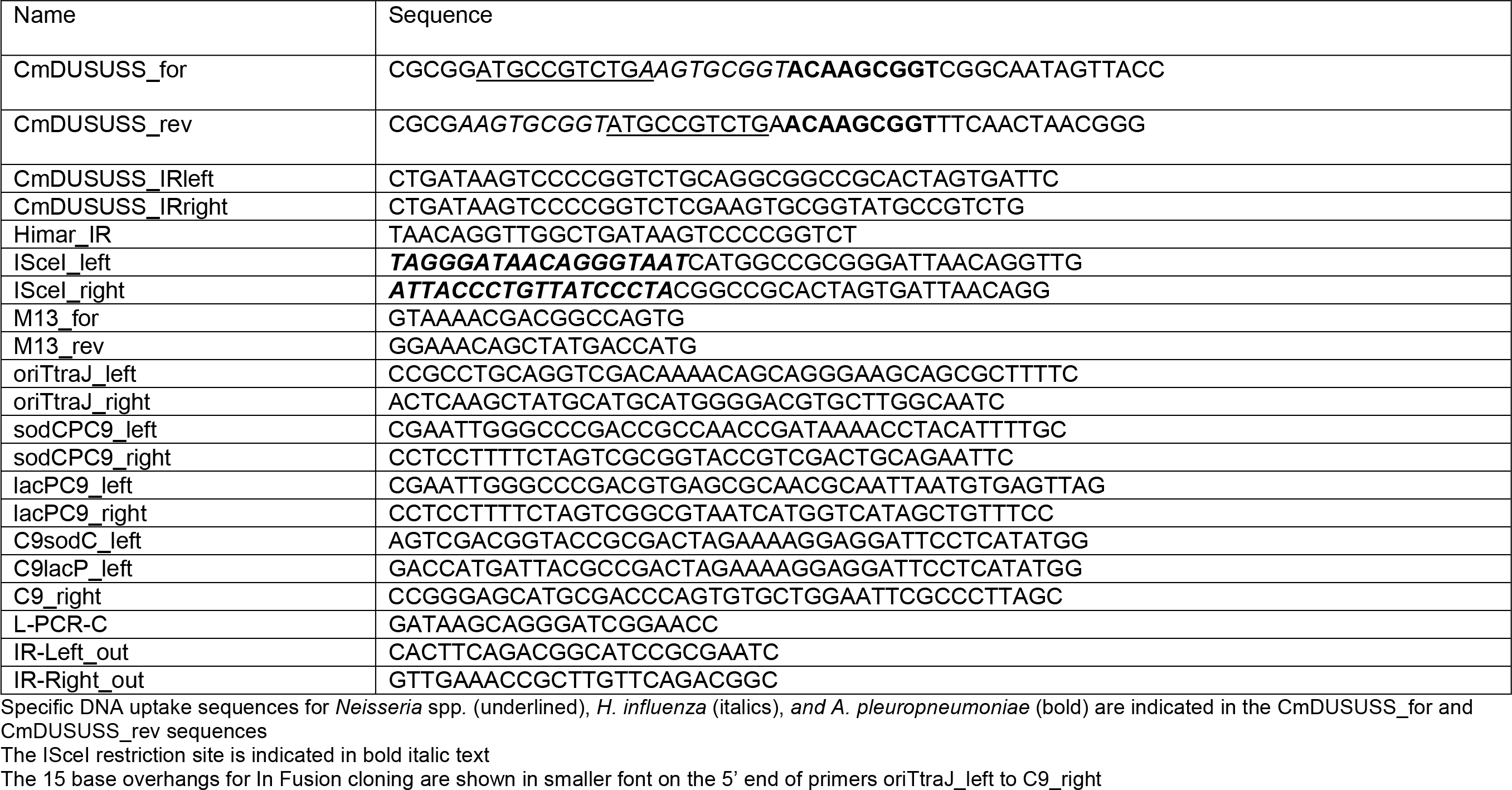
Primers used in this study.

**Table 2.** Location of *mariner* insertions in anaerobic mutants of *A. pleuropneumoniae*

| Gene | Product of disrupted gene | Gene location<sup>a</sup> | Location of insert<sup>a</sup> |
| --- | --- | --- | --- |
| *nhaB* | Na<sup>+</sup>/H<sup>+</sup> antiporter protein | 363852..365393C<sup>b</sup> | 364160 |
| *pflB* | Formate acetyltransferase | 1178587..1180899C | 1179054 |
|  |  |  | 1179780 |
|  |  |  | 1180383 |
| *atpC* | ATP synthase epsilon chain | 1922342..1922761C | 1922665 |
| *atpG* | ATP synthase gamma chain | 1924188..1925054C | 1924771 |
|  |  |  | 1924970<sup>c</sup> |
| *atpA* | ATP synthase alpha chain | 1925077..192661C | 1925488 |
|  |  |  | 1925791 |
|  |  |  | 1925826<sup>c</sup> |
|  |  |  | 1926363 |
| *atpB* | ATP synthase A chain | 1928081..1928869C | 1928204<sup>c</sup> |
|  |  |  | 1928575<sup>c</sup> |
| *fumC* | Fumarate hydratase | 2051573..2052967C | 2052243 |
| MIDG2331_02098 | D-alanyl-D-alanine carboxypeptidase | 2163589..2164257C | 2163863 |

<sup>a</sup>Location in the complete genome of MIDG2331 (accession number LN908249)
<sup>b</sup>Start and end positions of genes are indicated; those on the complementary strand are shown with the suffix C
<sup>c</sup>Insertions at these locations were mapped in 2 mutants each

**Table 3.** Location of *mariner* insertions in anaerobic mutants of *P. multocida* MIDG3277

| Gene | Product of disrupted gene | Gene location<sup>a</sup> | Location of insert<sup>a</sup> |
| --- | --- | --- | --- |
| MIDG3277_1258 | predicted glycosyl transferase | contig26:104818..105990<sup>b</sup> | contig26:105340 |
| *aroG* | deoxy-7-phosphoheptulonate synthase | contig17:68162..69247 | contig17:68149<sup>c</sup> |
| intergenic |  |  | contig37:10847<sup>d</sup> |
| MIDG3277_1016 | Hypothetical protein<sup>e</sup> | contig21:76828..77169C | contig21:77129 |
| *aroA* | 3-phosphoshikimate 1-carboxyvinyltransferase | contig21:79640..80962 | contig21:80029 |
|  |  |  | contig21:80287 |
| *sucC* | Succinate--CoA ligase subunit beta | contig3:86080..87246 | contig3:86908 |
| *ccmD* | Heme exporter protein CcmD | contig1:16084..16287 | contig1:16136 |
| *atpI* | F0F1 ATP synthase subunit I | contig38:36319..36726 | contig38:36262<sup>f</sup> |
| *atpG* | F0F1ATP synthase subunit gamma | contig38:40520..41389 | contig38:40497<sup>g</sup> |
| MIDG3277_0379 | Hypothetical protein | contig6:168..512<sup>h</sup> | contig6:434 |
| *pfeA* | Ferric enterobactin receptor | contig32:62820..64826C | contig32:64243 |
| *menA* | 1,4-dihydroxy-2-naphthoate octaprenyltransferase | contig28:97896..98801C | contig28:98875 |
| *menB* | 1,4-dihydroxy-2-naphthoyl-CoA synthase | contig28:34911..35768C | contig28:35383 |
| *menC* | O-succinylbenzoate synthase | contig28:33645..34658C | contig28:34159 |

<sup>a</sup>Location in the draft genome of MIDG3277 (accession number ERZ681052)
<sup>b</sup>Start and end positions of genes are indicated; those on the complementary strand are shown with the suffix C
<sup>c</sup>Insertion located 13 bp upstream of the start of *aroG*; insertion likely disrupts transcription of *aroG*
<sup>d</sup>Insertion located 75 bp upstream of the start of *degQ* and 147 bp upstream of the start of MIDG3277\_0890 encoding a putative NAD(P)H nitroreductase; insertion likely disrupts transcription of *degQ*
<sup>e</sup>This protein shares 100% identity with PM0836 (Genbank accession AAK02920) which was reported to be in vivo expressed (49)
<sup>f</sup>Insertion located 57 bp upstream of the start of *atpI* (MIDG3277\_1814); insertion likely disrupts transcription of *atpI* and remaining genes in the *atp* operon
<sup>g</sup>Insertion located 23 bp upstream of *atpG*; insertion likely disrupts transcription of *atpG* and remaining genes in the *atp* operon
<sup>h</sup>This predicted CDS is on a small contig and may be part of a larger gene (100% identity with the last 104 AAs of a 818 AA hypothetical protein
P1062\_0208970; Genbank accession ESQ7176

## Discussion

The *mariner* transposon does not require host-specific factors for transposition and has a minimal requirement for TA dinucleotides as its target site, allowing for greater distribution in genomes compared to transposons such as Tn*5* and Tn*10*, which have “hot spots” for insertion. Tn*10* insertion at hotspots was a severe limitation in our previous *A. pleuropneumoniae* signature tagged mutagenesis study (4, 11). Given the relatively AT rich genomes of *Pasteurellaceae* species, *mariner* should allow for creation of saturating mutant libraries in *A. pleuropneumoniae* and *P. multocida*.

Preliminary (unpublished) investigations in our laboratory using the vector pMinihimarRB1 (19), a kind gift from Dr D. Saffarini, revealed that *mariner* is functional in *A. pleuropneumoniae*, however disadvantages including high background with kanamycin selection and retention of plasmid in initial transconjugants limited the usefulness of this vector for genome-wide analysis of fitness using TraDIS, a next generation sequencing method for mapping insertion sites (14). We therefore decided to construct a *mariner* delivery vector incorporating the following desired components: a stringent selection gene carried by the mini-transposon; presence of DNA uptake sequences flanking the selection gene to allow for easy transfer of mutations between different strains via natural transformation; the presence of paired ISceI restriction sites just outside of the mini-transposon element to allow for elimination of plasmid reads during TraDIS; sequences to allow mobilization of the vector from a conjugal donor strain; and the C9 hyperactive mutant of the *Himar1 mariner* transposase gene (20), under control of either a constitutive or inducible promoter, encoded adjacent to the mini-transposon element to ensure stability of insertions.

Two novel mobilizable *mariner* vectors, pTsodCPC9 and pTlacPC9, were developed in this study. Both contain *mariner* mini-transposon elements carry a Cm resistance gene, known to provide stringent selection of mutants in *A. pleuropneumoniae* (21), flanked by paired DNA uptake sequences for each of *A. pleuropneumoniae*, *H. influenzae* and *Neisseria* spp. (22, 23). Although initially designed for mutagenesis of *A. pleuropneumoniae*, all features included were designed to facilitate use in other Gram-negative bacteria. As it is often desirable to transfer mutations into wild-type strains to confirm the effect(s) of gene mutation, DNA uptake sequences were designed into the transposon in order to facilitate transfer of mutations by natural transformation in the species which selectively take up DNA containing these elements (22, 23).

Restriction barriers can prevent or decrease the efficiency of plasmid delivery by electroporation into various bacteria (24, 25), we therefore added the *oriT* transfer origin and the *traJ* gene from the broad host-range conjugative plasmid RP4 (26) to facilitate delivery of the vectors by conjugation. Successful conjugal transfer of both vectors into MIDG2331 confirmed that these sequences were sufficient to allow mobilization from the DAP-dependent *Escherichia coli* strain MFD*pir* (27) which carries loci coding for the complete RP4 conjugation machinery. It is likely that the presence of the *oriT* alone would have been sufficient for mobilization, however, as binding of TraJ to the *oriT* initiates formation of the transfer machinery (26), a copy of this gene was included in the vectors. Use of MFD*pir* facilitates conjugation into wild-type recipient bacteria, instead of antibiotic resistant derivatives, as selection of transconjugants is achieved on agar that does not contain DAP which is required for growth of the donor strain (27).

The hyperactive C9 mutant transposase gene, previously shown to enhance the efficiency of *mariner* transposition (20, 28) was placed under transcriptional control of either the *A. pleuropneumoniae sodC* promoter, which we have shown to be active under all conditions investigated so far, both in *A. pleuropneumoniae* as well as in other *Pasteurellaceae* species (17), or the *E. coli lac* promoter which allows induction by IPTG, and which is functional in a variety of bacteria including *A. pleuropneumoniae* (4). Although both resulting plasmids, pTsodCPC9 and pTlacPC9, were functional in *A. pleuropneumoniae*, the former generated a lower frequency of transposants in this species, and only the latter generated transposants in *P. multocida*. Furthermore, IPTG induction of expression of the C9 transposase gene from the *lac* promoter resulted similar frequencies of transposition in both *A. pleuropneumoniae* and *P. multocida*, indicating it will likely be a useful genetic tool for other bacteria.

Using the pTlacPC9 vector, saturating libraries were generated for both *A. pleuropneumoniae* (insertions on average every 41 bp) and *P. multocida* (insertions on average every 17 bp), with insertions randomly distributed around the respective chromosomes. In both cases, pooled mutant libraries were prepared from initial selective plate cultures of the transconjugants (from multiple mating experiments) in order to limit selective expansion of clones. Under these conditions, extrachromosomal plasmid retained in the library can lead to high numbers of sequencing reads, interfering with TraDIS mapping. By engineering paired ISceI restriction sites (18 bp sequences not usually found in bacterial genomes) flanking the mini-transposon element in our vectors, we were able to effectively eliminate retained plasmid from the libraries prior to sequencing.

Both *A. pleuropneumoniae* and *P. multocida* are members of the *Pasteurellaceae* and are facultative anaerobes that infect the respiratory tracts of animals. Anaerobic growth is known to contribute to virulence of *A. pleuropneumoniae* (29). However, little is known about the importance of genes contributing to anaerobic growth in *P. multocida*. In this study, we have screened a limited number of *mariner* mutants of both pathogens to identify genes important for anaerobic growth in order to further validate the randomness of insertions in these libraries and usefulness of our approach. Of the 19 *A. pleuropneumoniae* mutants sequenced, 15 unique insertions in 8 different genes were identified (Table 2). Of 15 *P. multocida* mutants sequenced, all were in unique sites in the genome, with some insertions mapped to intergenic sites (Table 3). In both cases, mutants were selected from only four separate mating experiments, so it is possible that the four duplicate insertions mapped for the *A. pleuropneumoniae* mutants were due to clonal expansion following insertion, even though mutants were randomly selected from the primary counter selection plate. As clonal expansion can skew libraries for over representation of more rapidly growing mutants, steps should be taken to limit this prior to TraDIS analysis.

For both organisms, genes of the *atp* operon encoding the F_0_F_1_ ATP synthase appear to be essential for anaerobic growth. In *A. pleuropneumoniae*, 13 mutants had disruptions in four genes of the *atp* operon (nine unique sites). In *P. multocida*, two of the anaerobic mutants had insertions that mapped to sites upstream of genes in the *atp* operon (56 bp upstream of *atpI*, the first gene in the operon; and 24 bp upstream of *atpG*). These intergenic insertions likely disrupt transcription of all, or part, of the *atp* operon, and therefore production of functional F_0_F_1_ ATP synthase. STM studies have previously identified genes of this operon as important for *in vivo* survival of both *A. pleuropneumoniae* (4, 5), and of *P. multocida* (30), but the F_0_F_1_ ATP synthase has not previously been reported as essential for anaerobic growth of these bacteria. The number of unique *mariner* insertions mapping to the *atp* operons of these species in our current study provides strong evidence for the requirement of a functional F_0_F_1_ ATP synthase for their anaerobic growth. In the absence of oxidative phosphorylation, the F_0_F_1_ enzyme complex extrudes protons at the expense of ATP hydrolysis, to generate the driving force for solute transport and to maintain an acceptable intracellular pH value, and is required for function of certain enzymes involved in anaerobic respiration, such as formate hydrogenlyase (FHL) (31). Furthermore, activity of both FHL and F_0_F_1_-ATPase was found to facilitate the fermentative metabolism of glycerol in *E. coli* (32).

An insertion in *nhaB*, encoding a Na^+^/H^+^ antiporter, was also found to be important for anaerobic growth of *A. pleuropneumoniae* in our current screen. As with the *atp* operon, *nhaB* has been shown to be important for *in vivo* growth of *A. pleuropneumoniae* (33). The importance of interactions between proton and sodium cycles during both aerobic and anaerobic growth has been reported for certain pathogenic bacteria (34). The relative contributions of proton and sodium cycles to anaerobic growth of *A. pleuropneumoniae* and *P. multocida* warrant further investigation.

The importance of menaquinone biosynthesis for anaerobic growth of *P. multocida* was indicated by single insertions in each of *menA*, *menB*, and *menC*, as well as a single insertion three bp upstream of *aroG* (likely disrupting transcription), and two in *aroA*. Menaquinone is involved in anaerobic electron transport, and is derived from chorismate (pathway includes *menA*, *menB*, and *menC*), which in turn is derived from shikamate (pathway includes *aroA* and *aroG*).

Two *A. pleuropneumoniae* (*pflB* and *fumC*) genes identified encode proteins with known roles in anaerobic pathways. There were three separate insertions in *pflB* encoding pyruvate formate lyase which catalyses generation of formate via decarboxylation of puruvate. As mentioned above, the FHL complex is important in anaerobic respiration where it catalyses the conversion of formate into CO_2_ and H_2_ (35). A single insertion was mapped to *fumC* encoding fumarate hydratase, the enzyme responsible for catalysing the conversion of malate to fumarate. Fumarate has previously been shown to be an essential terminal electron acceptor during anaerobic respiration in *A. pleuropneumoniae* (29, 36).

Two of the *P. multocida* (*ccmD* and *sucC*) genes identified also have known anaerobic functions. Genes of the *ccm* operon are required for maturation of c-type cytochromes, including those involved in the electron transfer to terminal reductases of the anaerobic respiratory chain with nitrate, nitrite or TMAO (trimethylamine-N-oxide) as electron acceptors (37). The *sucC* gene encodes the beta subunit of succinate-CoA ligase, an enzyme in the aerobic citrate (TCA) cycle, where it catalyses hydrolysis of succinyl-CoA to succinate (coupled to the synthesis of either GTP or ATP). This enzyme also mediates the reverse reaction when required for anabolic metabolism, which can be particularly important under anaerobic conditions where the generation of succinyl-CoA via the oxidative pathway from 2-oxoglutarate is repressed (38, 39).

The remaining mutants in both *A. pleuropneumoniae* and *P. multocida* have insertions in (or upstream of) genes not directly linked with anaerobic growth, however further investigation is warranted to determine their possible contributions.

It is clear from our results that we have successfully constructed a *mariner* mini-transposon delivery vector capable of generating extremely large numbers of random mutants in *A. pleuropneumoniae*, *P. multocida*, and likely in other Gram-negative bacteria, which is amenable to genome-wide analysis of fitness using TraDIS. In preliminary experiments we have established that the pTlacPC9 vector can be used in *Aggregatibacter actinomycetemcomitans* and *Mannheimia haemolytica*, though it remains to be determined if the generated libraries are saturating.

The limited number of anaerobic-essential genes, identified individually via phenotypic analysis in the current study, mapped to different insertion sites, including some which had not previously been associated with anaerobic growth of *A. pleuropneumoniae* and *P. multocida.* This suggests that TraDIS analysis of the pooled *mariner* libraries subjected to the same screen will identify many more genes with functions contributing to anaerobic fitness. In future work, we will screen our *mariner* libraries under different *in vitro* and *in vivo* growth conditions, broadening our understanding of conditionally essential genes in the *Pasteurellaceae*.

## Materials and Methods

### Bacterial strains and culture

For generation of mariner transposon libraries, we chose two different *Pasteurellaceae* species, *A. pleuropneumoniae* and *P. multocida*. The *A. pleuropneumoniae* clinical serovar 8 isolate used, MIDG2331, was previously shown to be genetically tractable and has been fully sequenced (18). The *P. multocida* isolate was recovered from the respiratory tract of a calf in Scotland in 2008, and was shown to be sequence type 13, and part of clonal complex 13, in a multi-species multilocus sequence typing (MLST) study (40). This isolate, which has been labeled MIDG3277 in our collection, can be found in the pubMLST database (https://pubmlst.org/bigsdb?db=pubmlst_pmultocida_isolates) under the isolate name 22/4. The *Pasteurellaceae* isolates were routinely propagated at 37°C with 5% CO_2_ on Brain Heart Infusion (Difco) plates supplemented with 5% horse serum and 0.01% NAD (BHI-S-NAD). The *E. coli* strains used were: Stellar [*F-, ara,Δ(lac-proAB) [Φ 80d lacZΔM15], rpsL(str), thi, Δ(mrr-hsdRMS-mcrBC), ΔmcrA, dam, dcm*]; and MFD*pir* [MG1655 RP4-2-Tc::[*ΔMu1::aac*(3)*IV-ΔaphA-Δnic35-ΔMu2::zeo] ΔdapA::(erm-pir) ΔrecA*] (27). *E. coli* strains were maintained in Luria-Bertani (LB). Where appropriate, ampicillin (Amp; 100 μg/ml), chloramphenicol (Cm; 20 and 1 μg/ml for *E. coli* and *A. pleuropneumoniae*, respectively), and 0.3 mM diaminopimelic acid (DAP: required for growth of the MFD*pir* strain), were added to media.

### Genome sequencing, assembly and annotation

To obtain a draft genome sequence suitable for TraDIS analysis, genomic DNA was extracted from the *P. multocida* MIDG3277 strain using the FastDNA Spin kit (MP Biomedicals), and sequenced using an Illumina HiSeq 2000 at the Wellcome Sanger Institute. Illumina adapter sequences were trimmed from the reads using Cutadapt (41), and the trimmed reads were assembled using SPAdes (42). The assembled contigs were aligned to the complete genome sequence of *P. multocida* Pm70 (GenBank accession AE004439) using nucmer version 4.0.0 (43). Using the alignment, the contigs were reordered and reoriented to match the Pm70 reference genome, with one contig manually split where it overlapped the Pm70 origin. The reordered contigs were annotated using Prokka version 1.11 (44).

### Construction of the*mariner* mini-transposon vectors

All primers are listed in Table 1. CloneAmp HiFi PCR Premix (Takara) was used to amplify sequences for cloning, and the QIAGEN Fast Cycling PCR Kit (Qiagen) was used for verification of clones, using the respective manufacturer’s protocols. When required, blunt PCR products were A-Tailed using 5 U Taq polymerase and 0.2mM dATP prior to TA cloning into pGEM-T (Promega), according to the manufacturer’s protocol. Also, when required, DpnI digestion was used to remove plasmid DNA template from PCR products prior to cloning. All ligation products and In Fusion cloning products were transformed into *E. coli* Stellar cells (Clontech) by heat shock, according to the manufacturer’s protocol.

A *mariner* mini-transposon encoding Cm resistance was constructed in stages using pGEM-T as the vector backbone. Initially, the Cm cassette flanked by *A. pleuropneumoniae* uptake signal sequences (USS) was amplified from pUSScat (21) using primers CmDUSUSS_for and CmDUSUSS_rev, which further added DNA uptake sequences for *Neisseria* spp. and *H. influenzae* on both sides of the cassette. The resulting 956 bp amplicon was A-tailed and cloned into pGEM-T yielding the plasmid pTCmDUSUSS. The *mariner* inverted repeat (IR) sequence (TAACAGGTTGGCTGATAAGTCCCCGGTCT) was then added to either side of the CmDUSUSS cassette in two subsequent rounds of PCR amplification and cloning into pGEM-T. In the first round, primers CmDUSUSS_IRleft and CmDUSUSS_IRright added the last 19 bases of the *mariner* inverted repeat to either side of the cassette. The full IR sequence was then used as a primer (Himar_IR) to amplify the transposon cassette prior to cloning into pGEM-T to yield pTHimarCm. Finally, paired I-SceI restriction sites were added to either side of the transposon by PCR using primers ISceI_left and ISceI_right, and the product was cloned into pGEM-T to yield pTISceHimarCm. The full insert was sequenced using M13_for and M13_rev primers prior to further modifications of the plasmid.

All further cloning steps were performed using the In Fusion HD cloning kit (Clontech), according to the manufacturer’s protocol. A sequence containing the *oriT* and *traJ* gene was amplified from pBMK1 (45), a generous gift from Gerald Gerlach, using primers oriTtraJ_left and oriTtraJ_right, and cloned into NsiI/SalI cut pTISceHimarCm to yield pTISceHimarCmoriT. The C9 hyperactive *Himar1* transposase gene (20), amplified from pCAM45 (46) using primers C9_right and either C9sodC_left or C9lacP_left, was fused by overlap-extension (OE-PCR) PCR to either the *A. pleuropneumoniae sodC* promoter amplified from pMK-Express (17) using primers sodCPC9_left and sodCPC9_right, or to the *lac* promoter amplified from pBluescript II KS (Agilent Technologies) using primers lacPC9_left and lacPC9_right. The *sodCP*-C9 and *lacP*-C9 OE-PCR products were cloned into ZraI cut pTISceHimarCmoriT to yield the *Himar1 mariner* mini-transposon delivery vectors pTsodCPC9 and pTlacPC9, respectively. All inserts were confirmed by sequencing.

Purified pTsodCPC9 and pTlacPC9 plasmids were electroporated into the *E. coli* conjugal donor strain MFD*pir* (27), a generous gift from Jean-Marc Ghigo, with transformants recovered on LB containing 20 μg/ml Cm and 0.3 mM DAP.

### Bacterial mating and generation of *mariner* mutant libraries in *A. pleuropneumoniae* and *P. multocida*

Initially, the two different *mariner* mini-transposon delivery vectors, pTsodCPC9 and pTlacPC9, were evaluated for their ability to produce Cm-resistant mutants in *A. pleuropneumoniae* serovar 8 strain MIDG2331 following conjugal transfer from the DAP-dependent *E. coli* MFD*pir* donor strain. For mating experiments, donor and recipient bacteria were grown separately in broth culture to an optical density at 600 nm (OD_600_) of approximately 1.0 (cultures were adjusted to equivalent OD_600_). Two hundred microliters of recipient strain were mixed with 0 to 200 μl of donor strain (to give ratios of 1:1, 1:2, 1:4 and 1:8, as well as a control of recipient only). The bacteria were pelleted and re-suspended in 200 μl of 10 mM MgSO_4_, and 20 μl aliquots were spotted onto 0.45-μm nitrocellulose filters (Millipore) placed onto BHI-S-NAD agar supplemented with DAP and, when required for induction of the *lac* promoter, 1 mM IPTG (isopropyl-ß-D-galactopyranoside). Plates were incubated overnight at 37°C with 5% CO_2_, after which bacteria were recovered in 1 ml of sterile phosphate-buffered saline and aliquots were plated onto BHI-S-NAD agar supplemented with 1 μg/ml Cm. Selected transconjugants were tested, both before and after subculture on selective agar, by colony PCR for the presence of the Cm cassette using primers CmDUSUSS_for and CmDUSUSS_rev, and for presence of the plasmid backbone using primers oriTtraJ_left and oriTtraJ_right.

For construction of the *A. pleuropneumoniae mariner* library, a total of 14 matings were performed using the MFD*pir* + pTlacPC9 donor, with between 1400 and 6200 MIDG2331 transconjugants per mating resuspended in 3 ml BHI broth containing 50% (v/v) glycerol and stored in 1 ml aliquots at −80°C. A combined pooled library of transconjugants (56,664 insertion sites) was generated by mixing equal aliquots of each of the individual mutant pools. Southern blot and linker-PCR using AluI-digested DNA were performed as previously described (24, 47) in order to confirm single random insertion of the transposon in selected transconjugants.

The pTlacPC9 vector was further assessed for the ability to generate Cm-resistant mutants in the related bacterium, *P. multocida.* In total, 9 separate matings were performed using the MFD*pir* + pTlacPC9 donor and *P. multocida* MIDG3277 recipient, with between 2388 and 20,640 transconjugants selected per mating. Mutants were stored at −80°C as separate pools, and a combined pool (147,613 insertion sites), in 1 ml aliquots in BHI broth containing 50% (v/v) glycerol, as above.

### TraDIS analysis of the *A. pleuropneumoniae* and *P. multocida mariner* libraries

Genomic DNA was extracted from the pooled libraries of mutants using the FastDNA Spin kit (MP Biomedicals), according to the manufacturer’s protocol for bacterial cells. To assess the distribution of insertions in the pools prior to TraDIS, linker-PCR was performed as previously described (24) with the exception that, for PCR amplification, primer L-PCR-C was paired with either IR_left_out or IR_right_out (see Table 1) in place of L-PCR-L or L-PCR-R, respectively. For comparison, 2.5 μg aliquots of genomic DNA that were either untreated, or digested with ISceI to remove reads from residual plasmid, were used for linker PCR amplification of the left flank sequences, and the products were separated on 2% Nusieve agarose. Subsequently, 2 μg of ISceI-digested DNA was used to prepare Illumina libraries for TraDIS analysis, as previously described (25).

The TraDIS libraries were sequenced on an Illumina HiSeq 2500 at the Wellcome Sanger Institute. For the *A. pleuropneumoniae* library, TraDIS reads were mapped to the closed whole genome sequence of MIDG2331 (accession number LN908249). The *P. multocida* TraDIS reads were mapped to the draft genome of MIDG3277, constructed as described above. Reads were mapped using BWA mem (48), with an increased penalty for 5’ clipping (-L 100,5). To reduce background noise, aligned reads which did not match a TA site at the 5’ end of the alignment were excluded from further analysis.

### Preliminary screen for anaerobic mutants

Approximately 5000 mutants each for *A. pleuropneumoniae* and *P. multocida* (from 4 separate mutant pools) were screened for insertions in genes required for survival during anaerobic growth. The mutant pools were plated on BHI-S-NAD supplemented with 1 μg/ml Cm at a density of 75 to 150 colonies per plate. Following overnight incubation at 37°C with 5% CO_2_, colonies were transferred by replica plating onto 2 fresh selective plates. One plate was incubated with 5% CO_2_, and the other was placed in an anaerobic jar, at 37°C overnight. Mutants that failed to grow anaerobically were re-tested to confirm the growth defect, and the site of transposon insertion was determined for each by direct sequencing of linker-PCR products.

### Nucleotide sequence accession numbers

The complete sequences of the two *mariner* delivery vectors, pTsodCPC9 and pTlacPC9, have been deposited in Genbank under accession numbers MH644834 and MH644835, respectively. The draft genome sequence of the porcine clinical respiratory isolate of *P. multocida*, MIDG3277, has been deposited in the European Nucleotide Archive (ENA) under the accession number ERZ681052, with the raw reads available under the accession number ERR200085. The raw TraDIS reads for *A. pleuropneumoniae* are available in ENA under the accession number ERR271132. The raw TraDIS reads for *P. multocida* are available in ENA under the accession numbers ERR744003, ERR744016, ERR752316, ERR752329, ERR755725 and ERR755738.

## Acknowledgements

This work was supported by a Longer and Larger (LoLa) grant from the Biotechnology and Biological Sciences Research Council (BBSRC; grant numbers BB/G020744/1, BB/G019177/1, BB/G019274/1 and BB/G018553/1), the UK Department for Environment, Food and Rural Affairs and Zoetis awarded to the Bacterial Respiratory Diseases of Pigs-1 Technology (BRaDP1T) consortium.

Funding for LZ was provided by the BBSRC (grant number BB/C508193/1). MTGH was supported by the Wellcome Trust (grant number 098051). The funders had no role in study design, data collection and analysis, decision to publish, or preparation of the manuscript. The BRaDP1T Consortium comprises: Duncan J. Maskell, Alexander W. (Dan) Tucker, Sarah E. Peters, Lucy A. Weinert, Jinhong (Tracy) Wang, Shi-Lu Luan, Roy R. Chaudhuri (University of Cambridge); Andrew N. Rycroft, Gareth A. Maglennon, Jessica Beddow (Royal Veterinary College); Brendan W. Wren, Jon Cuccui, Vanessa S. Terra (London School of Hygiene and Tropical Medicine); and Janine T. Bossé, Yanwen Li and Paul R. Langford (Imperial College London).

